# Validation of a Demography-Based Adaptive QTc Formula using Pediatric and Adult Datasets Acquired from Humans and Guinea Pigs

**DOI:** 10.1101/2024.07.10.602984

**Authors:** Kazi T. Haq, Kate McLean, Grace C. Anderson-Barker, Charles I. Berul, Michael J. Shattock, Nikki Gillum Posnack

**Author notes:** Corresponding author: Nikki Gillum Posnack, Ph.D. Sheikh Zayed Institute for Pediatric Surgical Innovation Children’s National Hospital 111 Michigan Avenue, NW Washington, DC, USA 20010.

## Abstract

**Introduction:** A variety of QT rate-correction (QTc) formulae have been utilized for both clinical and research purposes. However, these formulae are not universally effective, likely due to significant influences of demographic diversity on the QT-HR relationship. To address this limitation, we proposed an adaptive QTc (QTcAd) formula that adjusts to subject demographics (i.e., age). Further, we compared the efficacy and accuracy of the QTcAd formula to other widely used alternatives.

**Method:** Using age as a demographic parameter, we tested the QTcAd formula across diverse age groups with different heart rates (HR) in both humans and guinea pigs. Utilizing retrospective human (n=1360) and guinea pig electrocardiogram (ECG) data from in-vivo (n=55) and ex-vivo (n=66) settings, we evaluated the formula’s effectiveness. Linear regression fit parameters of HR-QTc (slope and R²) were utilized for performance assessment. To evaluate the accuracy of the predicted QTc, we acquired epicardial electrical and optical voltage data from Langendorff-perfused guinea pig hearts.

**Results:** In both human subjects and guinea pigs, the QTcAd formula consistently outperformed other formulae across all age groups. For instance, in a 20-year-old human group (n=300), the QTcAd formula successfully nullified the inverse HR-QT relationship (R²=5.1E-09, slope=-3.5E-05), while the Bazett formula (QTcB) failed to achieve comparable effectiveness (R²= 0.20, slope=0.91). Moreover, the QTcAd formula exhibited better accuracy than the age-specific Benatar formula (QTcBe), which overcorrected QTc (1-week human QT: 263.8±14.8 ms, QTcAd: 263.8±7.3 ms, p=0.62; QTcBe: 422.5±7.3 ms, p<0.0001). The optically measured pseudo-QT interval (143±22.5 ms, n=44) was better approximated by QTcAd (180.6±17.0 ms) compared to all other formulae. Furthermore, we demonstrated that the QTcAd formula was not inferior to individual-specific QTc formulae.

**Conclusion:** The demography-based QTcAd formula showed superior performance across human and guinea pig age groups, which may enhance the efficacy of QTc for cardiovascular disease diagnosis, risk stratification, and drug safety testing.

**What is known:** Corrected QT (QTc) is a well-known ECG biomarker for cardiovascular disease risk stratification and drug safety testing. Various QT rate-correction formulae have been developed, but these formulae do not perform consistently across diverse datasets (e.g., sex, age, disease, species).

**What the study adds:** We introduce a novel QTc formula (QTcAd) that adapts to demographic variability, as the parameters can be modified based on the characteristics of the study population. The formula (QTcAd = QT + (|m|*(HR-HR_mean_)) – includes the absolute slope (m) of the linear regression of QT and heart rate (HR) and the mean HR of the population (HR_mean_) as population characteristics parametersˍUsing datasets from both pediatric and adult human subjects and an animal model, we demonstrate that the QTcAd formula is more effective at eliminating the QT-HR inverse relationship, as compared to other commonly used correction formulae.

## Introduction

The 12-lead electrocardiogram (ECG) is a widely used non-invasive diagnostic tool, with ECG parameters serving as cardiovascular disease biomarkers. However, studies in healthy humans^1,2^ and animals^3,4^ have reported significant rate-and age-dependent variations in baseline ECG metrics. These findings suggest that age is an independent demographic variable that modifies the electrical properties of the heart.

One of the most frequently used clinical ECG biomarkers to assess cardiovascular disease is the QT interval^5^, which is calculated as the interval between the onset of the QRS wave (marking the onset of ventricular activation) to the end of the T wave (marking the end of ventricular repolarization). However, the ventricular action potential duration, and hence the QT interval, has an inverse relationship with heart rate (HR) to physiologically allow for shortening or lengthening of the refractory period at fast or slow heart rates^6^. Accordingly, the effect of HR on the QT interval can complicate comparisons between subjects with different heart rates. To address this limitation, a correction formula was proposed to better predict QT at a resting HR, also known as the Bazett formula^7^, which provide a commonly utilized rate-adjusted QT that is referred to as a corrected QT (QTc).

Despite its widespread use in clinical studies and practices, the Bazett correction formula (QTcB) underperforms when applied to individuals with both fast and slow HR^8,9^. To enhance performance, a number of alternative universal QT correction formulae have been proposed^10^, including Framingham (QTcFram)^11^, Fridericia (QTcF)^12^, and Hodges (QTcH)^13^ formulae. The QTcB formula has also been shown to be less effective in preclinical animal models, which prompted the development of alternative species-specific formulae for use in rats^14^, dogs^15^, and monkeys^16^.

There are conflicting reports on the performance of various QTc formulae, which is likely influenced by the population characteristics within each study. For example, QTcB showed superiority in patients with congenital long QT syndrome (LQTS)^17^, but it was found to be inferior to both QTcF and QTcFram formulae in another study^6^. While another study by Luo et. al suggested that the QTcH formula outperformed QTcB, QTcF, and QTcFram formulae in adult patients with normal ECGs^18^. Similarly, in preclinical models, a rodent-specific formula was noted as superior in one study^14^, but failed to demonstrate satisfactory rate-correction in another study – in which it worsened the QT-HR inverse relationship^19^. Given the limitations with creating a “universal” QTc formula, some studies^9,20^ have suggested that inter-subject variability is too great within a given population, and that individual-specific QTc formulae should be adopted for improved accuracy.

It is apparent that demographic characteristics within a given study population can contribute to variable ECG metrics, which can pose significant limitations when applying a single, universal QTc formula to an entire dataset. To address this limitation, we developed a demography-based universal QT correction formula, which we termed an “adaptive QTc formula (QTcAd).” Given the known age-dependent effects on HR and QT, we tested the performance of QTcAd using age as a demographic variable. To test performance, we utilized retrospective ECG data acquired from multiple age groups within both human and guinea pig subjects. We compared the performance of the QTcAd formula to other widely used formulae, including the age-specific Benatar QTc formula (QTcBe)^24^. Further, we employed optically-acquired voltage data from Langendorff-perfused guinea pig hearts to validate whether the QTc values closely approximated the activation-repolarization time. We also compared the performance of a population-based adaptive formula (QTcAd) with an individual-based adaptive formula using an ex-vivo guinea pig ECG dataset, in which intra-subject HR was directly modulated by external pacing and a pharmacological agent. Finally, we demonstrate that the choice of QT correction formula has a direct impact on data interpretation.

## Material and Methods

### Study population

#### Animals

This study includes guinea pigs from two centers: Children’s National Research Institute (CNRI) in Washington, DC, United States, and the British Heart Foundation Centre of Research Excellence at King’s College London (KCL), United Kingdom. Animal procedures were approved by the Institutional Animal Care and Use Committee within CNRI, in accordance with the guidelines outlined in the National Institutes of Health’s Guide for the Care and Use of Laboratory Animals, as well as the Animal (Scientific Procedures) Act (1986) under the United Kingdom Home Office licenses PPL 70/7491 and PPL PA527CE7C, with ethical approval from the KCL Animal Welfare and Ethical Review Body.

Animals at CNRI were mixed-gender Dunkin-Hartley guinea pigs (n=66) sourced from Hilltop Lab Animals in Pennsylvania, USA and Charles River Laboratories in Quebec, Canada. Animals at KCL (n=8) were adult male Dunkin-Harley guinea pigs sourced from Marshal Bioresources in East Yorkshire, UK. At both centers, animals were accommodated in traditional acrylic cages within the research animal facility, where they were subjected to standard environmental settings. These included a 12-hour light/dark cycle, temperatures maintained between 18 and 25°C, and humidity levels ranging from 30% to 70%. At CNRI, animals were divided into three age groups: neonates (1-2 days old), younger adults (1-3 months old), and older adults (18-22 months old). At KCL, only adult male Dunkin Hartley guinea pigs were used (300-350g). Animals were group-housed, when possible, with free access to food and drinking water at all times. **Supplemental Table 1** summarizes the number of animals used for *in-vivo* and *ex-vivo* ECG recordings, and optical imaging.

#### Human subjects

In this retrospective IRB approved study (#12146), the study population was derived from data collected at Children’s National Hospital. Standard 12-lead ECG data were collected from patients admitted between 1993 and 2015.

Only patients meeting the following criteria were included:

I. Individuals with confirmed normal ECG readings, as assessed by the consulting physician.
II. For individuals with multiple ECG recordings, subjects were included only if all prior ECG readings were confirmed to be normal.

ECG data were collected using the MUSE Cardiology Information System (GE Medical Systems, Menomonee Falls, WI), a standard system for recording and analyzing 12-lead ECGs. ECGs were recorded using the MAC 5500 ECG system of GE Healthcare, operating at a paper speed of 25 mm/s, an amplitude of 10 mm/mV, and a sampling rate of 250 Hz. A total of 1360 individual ECG recordings were utilized for the study, which were grouped into 8 age categories as shown in **Supplemental Table 2**. It is important to note that each group consisted of individuals with the exact same age, which reduces intersubject variability compared to studies that utilize a wider age range. The study was conducted in compliance with ethical standards and received approval from the institutional review board of Children’s National Hospital.

### *In vivo* and *ex vivo* ECG recordings in guinea pigs

*In-vivo* ECG recordings in the CNRI cohort were performed as previously described^25^. Briefly, animals were anesthetized with 3–4% isoflurane for 10 minutes. Subcutaneous electrodes were used to collect ECG signals (lead I) at a sampling frequency of 1000 Hz using a PowerLab acquisition system (ADInstruments, Colorado Springs, CO). To obtain *ex-vivo* pseudo-ECGs, guinea pigs were anesthetized and the heart was excised as previously described^3^. After cannulation of the aorta, the intact heart was transferred to a temperature-controlled (37°C), constant pressure (68–70 mm Hg) Langendorff-perfusion system. Excised hearts were perfused with a modified Krebs–Henseleit buffer solution, comprising (in mM) 118.0 NaCl, 3.3 KCl, 1.2 MgSO_4_, 1.2 KH_2_PO_4_, 24.0 NaHCO_3_, 10.0 glucose, 2.0 sodium pyruvate, 10.0 HEPES buffer, and 2.0 CaCl_2_, bubbled with carbogen. Isolated hearts were allowed to equilibrate for 10–15 minutes on the Langendorff-perfusion system, followed by recording of pseudo-ECGs in a lead I configuration. Signal processing for both *in-vivo* and *ex-vivo* ECG recordings was performed using filters and amplification via a differential amplifier (high-pass filter (10 Hz), low-pass filter (100Hz), and gain amplifier, along with a 60 Hz noise eliminator (Hum Bug: Digitimer, Fort Lauderdale, FL). ECG metrics were determined by averaging 10 consecutive beats using LabChart 8 software (ADInstruments, Colorado Springs, CO).

### Heart rate adjustment protocol

*Ex-vivo* ECG recordings at KCL were obtained using an isolated working heart system, perfused with a modified buffer solution (perfusate (in mM): 114.0 NaCl, 4.0 KCl, 1.0 MgSO_4_, 1.1 NaH_2_PO_4_, 24.0 NaHCO_3_, 11.0 glucose, 2.0 sodium pyruvate, and 1.5 CaCl_2_). Briefly, hearts were cannulated and stabilized in Langendorff-mode for 15 minutes. Subsequently, they were switched to working mode and allowed an additional 15 minutes for stabilization. After equilibration, the pseudo-ECG was captured using silver wire electrodes arranged in a lead II configuration and recorded via a PowerLab acquisition system. A gradual reduction in HR was performed through the administration of incremental doses of ivabradine added to the perfusate buffer, starting at 0.2 µM and reaching a maximum of 0.6 µM, to achieve a target HR of approximately 120 bpm. HR was then incrementally increased through atrial pacing in 20 bpm increments (with 2 minutes at each pacing rate), until reaching a maximum HR of 310 bpm. This protocol facilitated data collection across a range of physiological heart rates to obtain individual-specific ECG datasets.

### QT interval approximation by optical mapping

To validate the corrected QT in guinea pigs, optically-acquired voltage signals were recorded from the epicardial surface of excised, intact heart preparations as previously described^3^. Briefly, after *ex-vivo* ECG recordings were performed, 12 μM (-/-) blebbistatin was introduced to reduce contractile function and mitigate motion artifacts in optically-acquired signals. Subsequently, the hearts were loaded with a potentiometric dye (RH237) and an LED light source (535±25 nm) was used to illuminate the epicardial surface. Fluorescence signals (>700 nm) were acquired using an Optical Mapping System (Mapping Labs LtD, Oxford UK). The optically approximated QT (QTc_op) interval at the resting HR was determined by calculating the duration between the breakthrough time (the onset of activation) and the time when the signal returns to baseline (indicating repolarization completion) from epicardial voltage signals.

### Formulation of adaptive QTc formula

We devised a generalized QT correction formula inspired by linear QT correction formulae from previous studies^11,13,24^. However, employing fixed parameters in these formulae restricts their applicability across diverse populations. This variability stems from differences in population-specific demographics that influence the QT and HR relationship. To address this limitation, we developed a more flexible expression for QT correction based on fundamental linear regression principles (**Supplemental Methods, Supplemental Figure 1**). This allows for adjustments in the formula parameters according to the characteristics of a given dataset. The "adaptive" QTc formula is represented as follows:

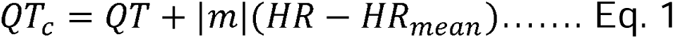

where, *m* is defined as the absolute value of the slope of the linear regression of QT and HR and

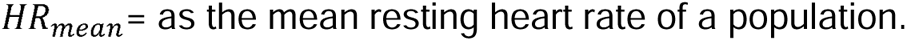

### Estimation of general parameters for different data sets and demographics

In Eq. 1, the values of m and HR*_mean_* can be estimated from uncorrected QT and HR values from a specific dataset. However, the application of the formula given by Eq. 1 requires such ECG datasets to be available for parameter estimation. To overcome this limitation, specifically in relation to age, we have developed an age-based regression model using normal human ECG data, which provides an estimation of parameters for the formula in Eq. 1 (**Figure 1A,B**) if the age of the subject is known. Similar models could be generated for other demographic variables.

**Figure 1.**
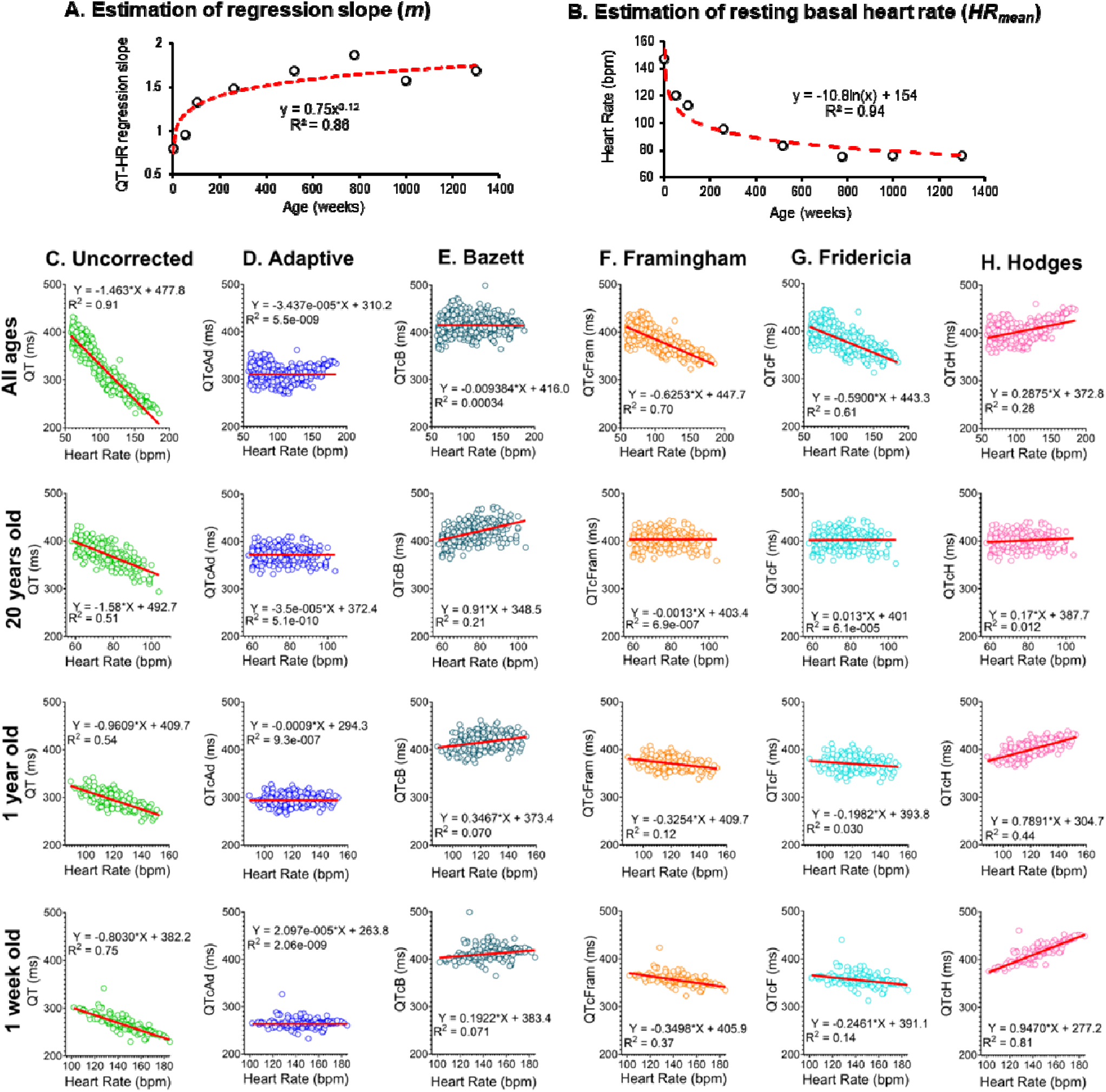
Estimation of human age-specific QTcAd parameters and comparison with other QTc formulae. **A)** Regression model depicted uncorrected QT-HR regression slope (|m|) with respect to patient age. Mean values of the linear regression slope were obtained from 12-lead ECG recordings from n=1360 individuals, across eight different age groups. **B)** Regression model illustrating the relationship between mean resting HR and age, using the same subjects. **C)** Linear regression fit of the uncorrected QT-HR relationship in healthy individuals (n=900), including 1-week old (n=300), 1 year old (n=300), and 20 years old (n=300). **D-H)** Linear regression fits of QTc-HR using different correction formulae in the same subjects. Corresponding equations and goodness of fit (R^2^) are provided. QT: Uncorrected QT, QTcAd: Adaptive QTc, QTcB: Bazett QTc, QTcFram: Framingham QTc, QTcF: Fridericia QTc, QTcH: Hodges QTc, bpm = beats per minute.

### Performance evaluation of adaptive formula

We assessed the performance of our adaptive formula using data collected from both human subjects and guinea pigs. To evaluate its efficacy in mitigating the inverse correlation between QT and HR, we compared our adaptive QTc formula against other established formulas, including:

Hodges: *QTcH - QT* + 1.75(*HR* - 60)

Bazett: *QTcB - QT/RR*^l/2^

Framingham: *QTcFram - QT* + 0.154(1 - *RR*)

Water: *QTcW - QT* - 0.087(*RR* - 1)

Fridericia: *QTcF - QT/RR*^l/3^

Benatar: *QTcBe = QT+α(1-RR)*

Wherein, HR is in bpm, RR is in seconds, and α is the slope of QT-RR regression line. The efficacy of each formula was assessed by examining the slope of the regression line and the degree of correlation, as indicated by the R^2^ value.

### Statistical Analysis

Descriptive statistics are presented as mean ± SD. Group comparisons were performed using 1-or 2-way ANOVA with Holm-Sidak correction for post-hoc analysis. Comparison between two groups were performed using either Student’s t-test or Wilcoxon test. Statistical significance was defined as p < 0.05, indicated in each figure by an asterisk. Regression analysis was employed to evaluate the relationship between variables, with the slope of the regression line and R^2^ value indicating both the direction and strength of correlation.

## Results

### Subject demographics

In the animal cohort, we conducted *in-vivo* and *ex-vivo* ECG recordings involving 66 guinea pigs. Animals displayed a trend of decreasing HR and increasing QT interval with advancing age (**Supplemental Table 1**). Statistical significance was predominantly observed when comparing neonates with older adult animals. For instance, in *ex-vivo* assessments, neonatal HR was significantly faster than that of adult animals (*Neonate*: 250.7±22.9 bpm, *Younger adult*: 227.5±35.2 bpm, *Older adult*: 207.7±34.6 bpm, p<0.0001). However, HR deceleration and QT prolongation in older animals did not reach statistical significance compared to their younger counterparts in either experimental group. In the KCL guinea pig cohort, all animals were young adults, weighing between 300-350g. For the ivabradine studies, the mean baseline HR and QT interval were 258±14.1 bpm and 139.5±5.9 ms, respectively.

A total of 1360 patients with normal ECG recordings were included in this retrospective analysis. The demographic distribution is summarized in **Supplemental Table 2**, with a comparable number of male (46.3%) and female (53.7%) patients. Analogous to observations in guinea pigs, both sexes exhibited age-dependent variations in HR and QT (**Supplemental Table 2**). For instance, HR decreased significantly with age in males (1 week: 147.3±16.1 bpm, 1 year: 119.9±11.8 bpm, 20 years: 75.1±10.5 bpm, p<0.0001), while the QT interval prolonged significantly (1 week: 263.3±14.2 ms, 1 year: 294.1±16.0 ms, 20 year: 367.6±20.0 ms, p<0.0001). Similar age-dependent adaptations were observed in females. Except for QT values in the 10-year and 20-year age group, HR and QT remained similar between male and female patients.

### Adaptive formula using “age” as a demographic variable

#### Estimation of parameters from population data

We utilized data from three specific age groups (1 week, 1 year, and 20 years) in humans, and from all age groups of the guinea pigs to illustrate the process of deriving the QT-HR regression line slope and mean resting HR for the QTcAd formula, assuming a dataset is available. Subsequently, we employed these same age groups to evaluate and compare the efficacy of various correction formulae. Upon examining the demographic profile of our study population (see **Supplemental Table 1 and 2**), notable variations were observed in HR and QT intervals across different age groups. We postulated that these demographic variances warranted unique QT correction formulas tailored to each subgroup. Consequently, by substituting the absolute linear regression slope (*m*) and mean resting HR (*HR_mean_*) values (**Table 1**) into Eq.1 for each age group, we derived a series of correction formulae. It is important to note that each adaptive formula corresponds to a specific age group and comprises a distinct set of parameters.

**Table 1.**
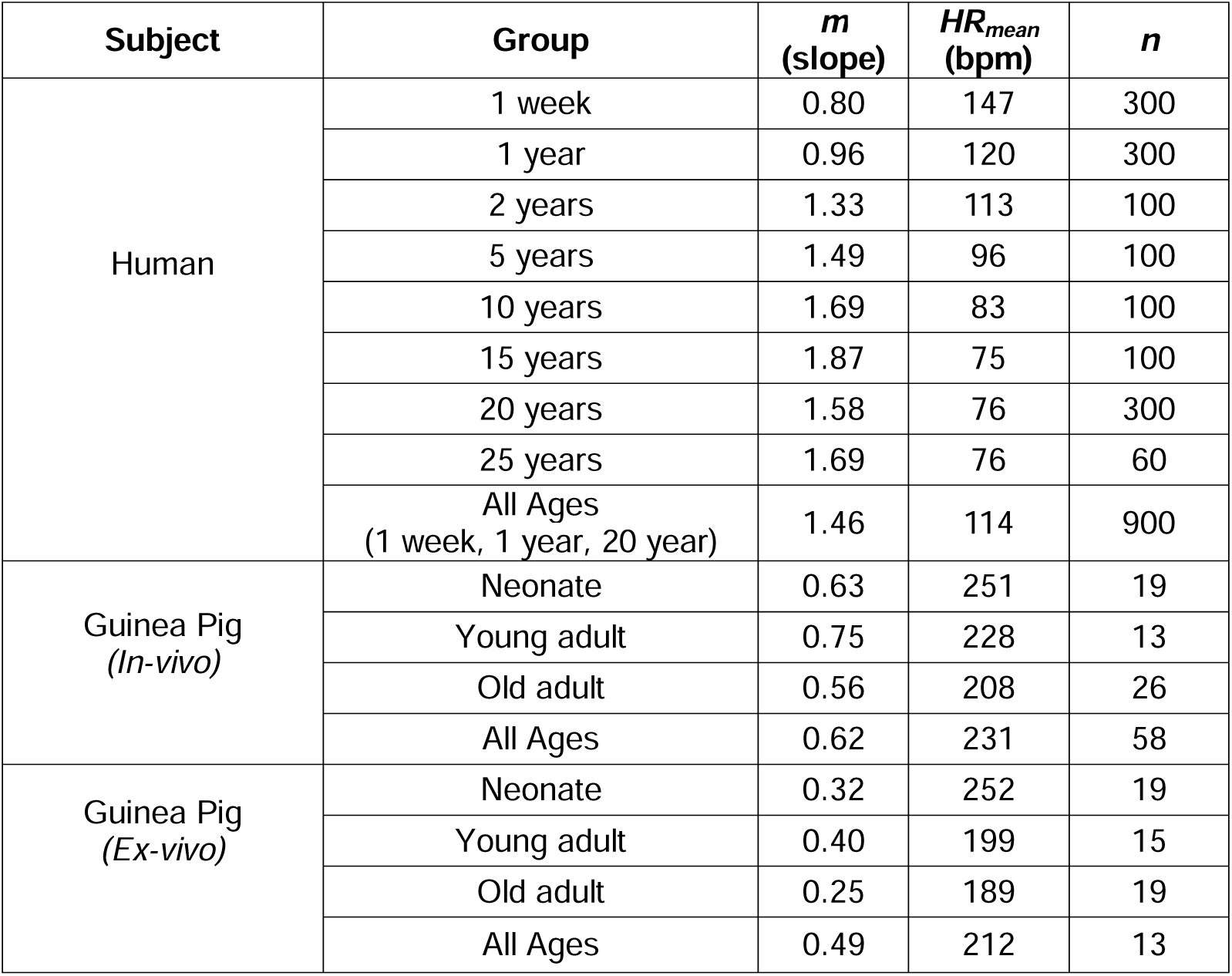
Age-specific QTcAd formula parameter values for specific age groups. The absolute value of QT-HR linear regression slope is denoted by *m* and the mean resting heart rate is denoted by *HR_mean_*.

| Subject | Group | $m$<br>(slope) | $HR_{mean}$<br>(bpm) | $n$ |
| --- | --- | --- | --- | --- |
| Human | 1 week | 0.80 | 147 | 300 |
|  | 1 year | 0.96 | 120 | 300 |
|  | 2 years | 1.33 | 113 | 100 |
|  | 5 years | 1.49 | 96 | 100 |
|  | 10 years | 1.69 | 83 | 100 |
|  | 15 years | 1.87 | 75 | 100 |
|  | 20 years | 1.58 | 76 | 300 |
|  | 25 years | 1.69 | 76 | 60 |
|  | All Ages<br>(1 week, 1 year, 20 year) | 1.46 | 114 | 900 |
| Guinea Pig<br>(*In-vivo*) | Neonate | 0.63 | 251 | 19 |
|  | Young adult | 0.75 | 228 | 13 |
|  | Old adult | 0.56 | 208 | 26 |
|  | All Ages | 0.62 | 231 | 58 |
| Guinea Pig<br>(*Ex-vivo*) | Neonate | 0.32 | 252 | 19 |
|  | Young adult | 0.40 | 199 | 15 |
|  | Old adult | 0.25 | 189 | 19 |
|  | All Ages | 0.49 | 212 | 13 |

#### Estimation of parameters from a regression model

We utilized data from eight young human age groups (n=1360) to construct an age-based regression model for estimating HR-QT linear regression slopes *(m)* and mean resting heart rate (HR*_mean_*) for each group. Both regression models exhibited a strong correlation (**Figure 1A,B**). Consequently, the regression fits facilitated the estimation of parameters for the QTcAd formula through the following relationships.

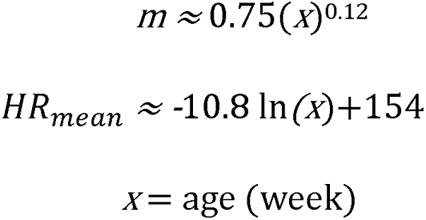

From Eq. 1,

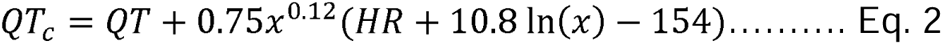

### QT correction formulae comparison using human subject datasets

We evaluated the performance of the QTc adaptive formula against several commonly used alternatives (**Figure 1C-H**). For this comparison, we utilized ECG data collected from four different groups (1 week old, 1 year old, 20 years old, and all ages combined). In human subjects, the QTcAd formula achieved nearly zero regression slope and demonstrated no correlation between HR and QTc in either the comprehensive age dataset or the age-specific datasets (**Figure 1D**, **Supplemental Table 3**). For example, in the comprehensive age dataset, the QTcAd formula resulted in a regression line slope of -3.4E-05 and an R^2^ value of 5.5E-09 (n=900 individual patients, **Figure 1D**). In comparison, other QTc formulae underperformed, including both the QTcFram (*m:* -0.63, R²: 0.7) and QTcF (*m:* - 0.59, R²: 0.6) equations (**Figure 1F,G**). The QTcAd formula also achieved a nearly zero regression slope for each of the tested age groups, including 1 week old neonates (*m:* 2.1E-05, R²: 2.1E-09, n=300), 1 year old infants (*m:* -0.0009, R²: 9.3E-07, n=300), and 20-year old adults (*m:* -3.5E-05 slope, R²: 5.1E-010, n=300; **Figure 1D**). However, the performance of other QTc formulae varied significantly across different age groups. As one example, the QTcB formula showed satisfactory performance in the comprehensive age dataset (*m:* 0.0094, R²: 0.0003, n=900), but performed more poorly in the 20-year old age group (*m:* 0.91, R^²^: 0.21, n=300; **Figure 1E**). Note that QTcF formula yielded satisfactory results in the 20-year age group (*m:* 0.013, R²: 6.1E-05), yet failed to achieve effectiveness in the comprehensive age dataset. Finally, the QTcH formula performed poorly in both of the younger age groups, including 1-year old (*m:* 0.79, R²: 0.44) and 1-week old (*m:* 0.95, R²: 0.81). A complete list of derived data for each of the tested QTc formulae is provided in **Supplemental Table 3.**

### QT correction formulae comparison using guinea pig datasets

To further assess the efficacy of the QTcAd formula, we tested its performance in a guinea pig dataset comprised of four different groups (neonate, younger adult, older adult, all-ages combined). In addition to the formulae employed in humans, we also compared the QTcAD formula against the Van de Water formula developed for dogs^26^. Similar to the human dataset, the QTcAd formula consistently produced a flat regression line and demonstrated no correlation between HR and QTc, both in vivo and ex vivo datasets (**Supplemental Figure 2,3, Supplemental Table 4-5**). For example, in the comprehensive age dataset, the QTcAd formula resulted in a regression line slope of 5.4E-06 and an R^2^ value of 2.4E- 010 for *in-vivo* measurements (**Figure 2B**) and a slope of 5.1E-08 and an R^2^ value of 1.3E-014 for *ex-vivo* measurements (**Figure 2E**). In comparison, the QTcB formula (*m:* -0.35, R²: 0.21, *in-vivo*) and other commonly employed QTc formulae consistently underperformed (**Figure 2C**, **Supplemental Table 4,5**). The QTcAd formula also achieved superior performance for each of the guinea pig age groups tested *in vivo*, including older adults (*m:* 1.56E-005, R²: 2E-009), younger adults (*m:* -2.62E-005, R²: 4.6E-009), and neonates (*m:* -2.04E-005, R²: 2.3E-009; **Figure 2B**). Similar results were obtained when the QTcAd formula was applied to datasets from *ex vivo*, intact Langendorff-perfused heart preparations in older adults (*m:* -9.1E-006, R²: 1.2E-010), younger adults (*m:* -4.63E-005, R²: 6.5E-009), and neonatal guinea pigs (*m:* -2.41E-005, R²: 2.4E-009; **Figure 2E**, **Supplemental Figure 3**).

**Figure 2.**
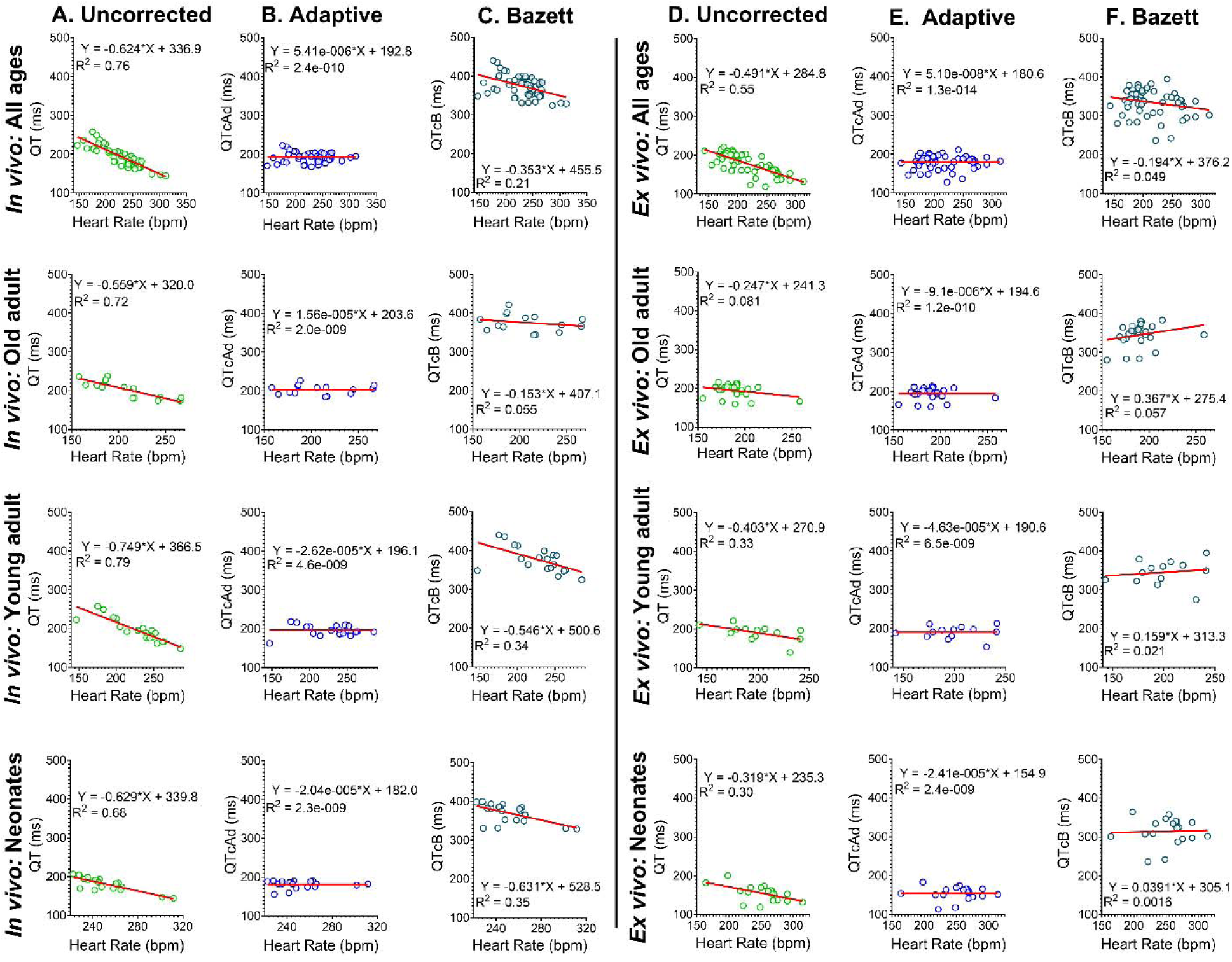
Comparison of QTc formulae across specific guinea pig age groups. **Left:** *In vivo* linear regression analysis of the QT-HR relationship in guinea pigs across 4 different age groups, including all ages, neonates (n=19), young adults (n=13), and old adults (n=26). **A)** uncorrected QT-HR, **B)** QTcAd-HR, **C)** QTcB-HR derived from the same animals and age groups. **Right:** *Ex vivo* linear regression analysis of the QT-HR relationship in guinea pigs across 4 different age groups, including all ages, neonates (n=21), young adults (n=19), and old adults (n=15). **D)** uncorrected QT-HR, **E)** QTcAd-HR, F) QTcB-HR derived from the same animals and age groups. Corresponding equations and goodness of fit (R2) are provided. QT: Uncorrected QT, QTcAd: Adaptive QTc, QTcB: Bazett QTc, bpm = beats per minute.

The performance of other formulae varied across age groups, when applied to both *in vivo* and *ex vivo* measurements. For example, the QTcB and QTcFram formulae exhibited satisfactory performance in *ex vivo* neonatal and older adult guinea pig datasets, respectively (QTcB: *m*: 0.039, R²: 0.0016; QTcFram: *m:* -0.077, R^2^: 0.0039; **Figure 2E**, **Supplemental Table 5**), but demonstrated poorer performance in the all-ages group (QTcB: *m:* -0.19, R²: 0.05; QTcF: *m:* -0.36, R²: 0.21; **Figure 2F**, **Supplemental Table 5**). The Van de Water formula (QTcW), developed for dogs, also underperformed in the all-age guinea pig group when applied to both *in vivo* (*m:* -0.51, R^2^: 0.68) and *ex vivo* datasets (*m:* -0.38, R^2^: 0.042). The effectiveness of each QTc formulae is shown in **Figure 3**, and the derived data values are reported in **Supplemental Tables 4,5.**

**Figure 3.**
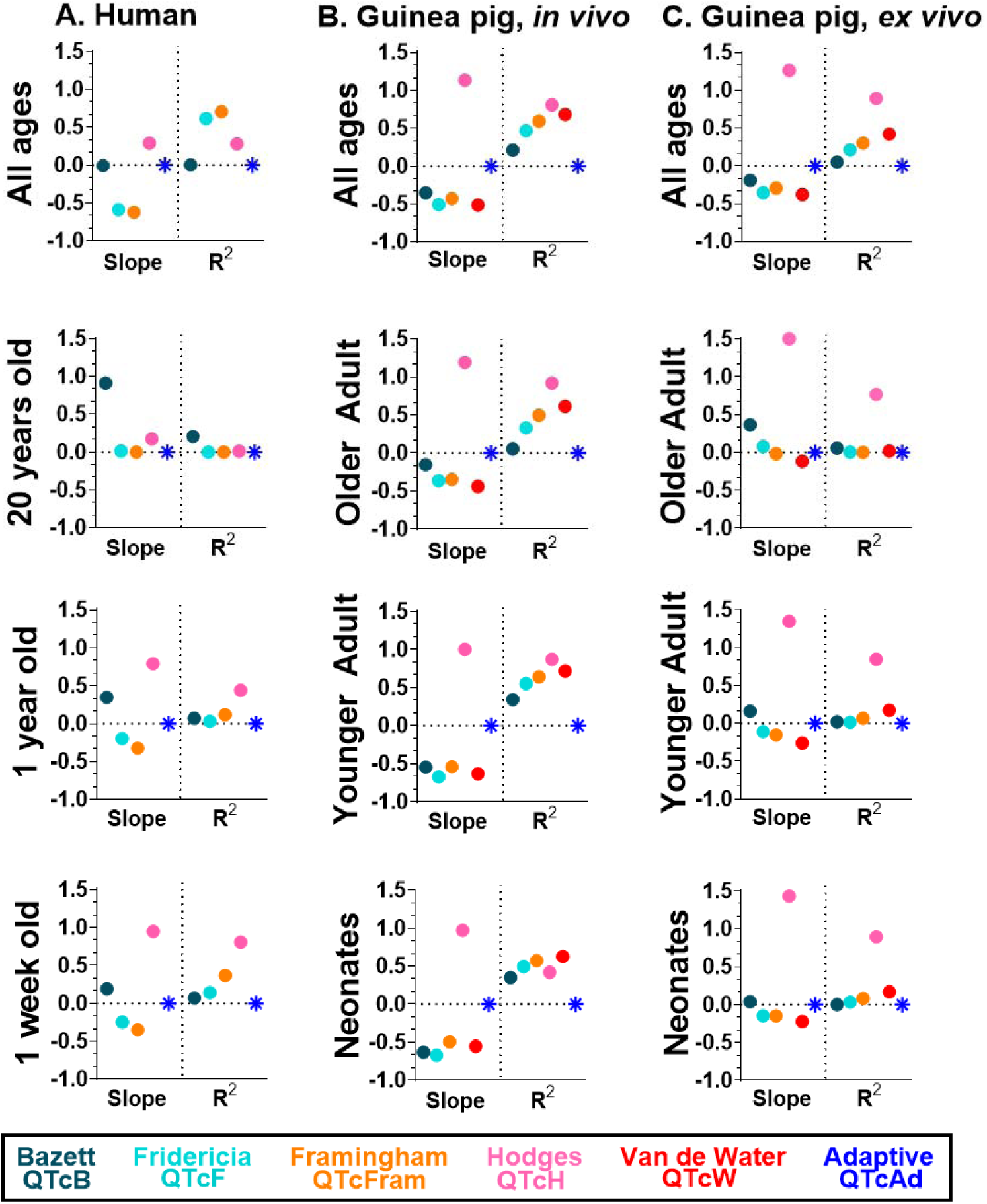
Comparison of QTc formulae efficacy in humans and guinea pigs. Comparison of QTc-HR linear regression slopes and goodness of fit (R2) values derived using different correction formulae. **A)** Dataset from healthy humans across four age groups, including all ages combined (n=900), 1-week (n=300), 1-year (n=300), and 20-year (n=300). **B)** Dataset from *in vivo* ECG recordings in healthy guinea pigs across four age groups, including all ages combined (n=58), neonates (n=19), young adults (n=13), and old adults (n=26). **C)** Dataset from *ex vivo* ECG recordings in healthy guinea pigs across four age groups, including all ages combined (n=42), neonates (n=21), young adults (n=19), and old adults (n=15).

### Evaluation of QT correction via optical mapping validation

Next, we tested whether the corrected QT intervals closely approximated the activation-repolarization time of the heart. For these studies, QTc was calculated from optically-acquired voltage data (QTc_op) from *ex vivo* heart preparations (**Figure 4A-C**) and the QTc_op values were compared to QTc values derived via different correction formulae. QTc_op values from individual guinea pig heart preparations are shown in **Figure 4D**, which aligned most closely with uncorrected QT and QTcAd values – compared to all other formulae. To quantify the offset of the approximation, we measured the difference between the uncorrected QT (from the pseudo-ECG recording) and the QTc_op values (ΔQT) – and then compared the measured duration difference between each correction formula (**Figure 4E**). The duration difference using the QTcAd formula (39.9±20.5 ms) was comparable to the duration difference using the QTc_op measurement (40±22.68 ms). All of the other tested correction formulae displayed a significant duration difference (QTcH > QTcB > QTcFram > QTcF > QTcW; n=44).

**Figure 4.**
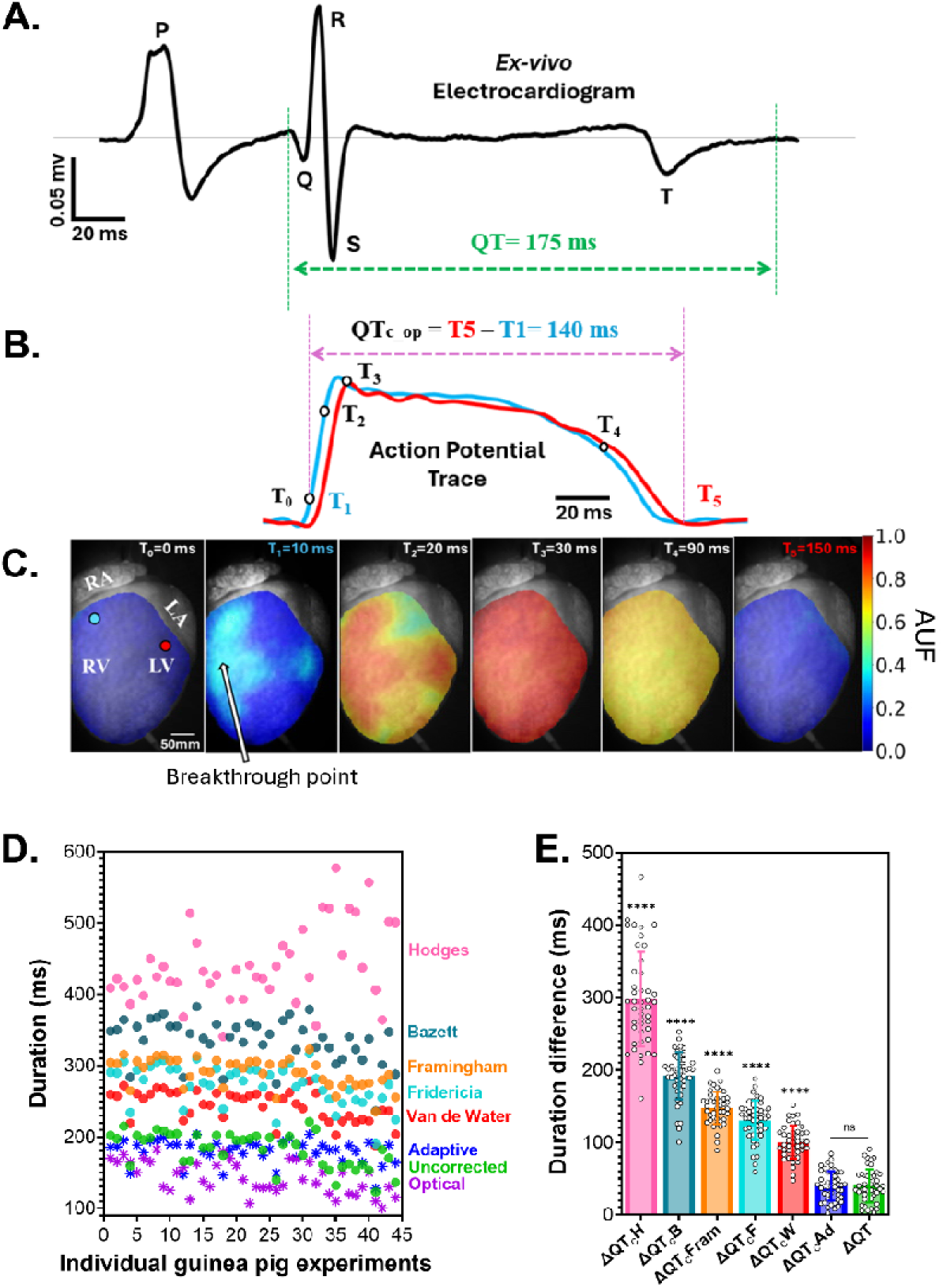
Evaluation of QT correction via optical mapping validation. **A)** Representative *ex vivo* pseudo-ECG trace from an excised guinea pig heart, with markers to show the uncorrected measurement. **B)** Representative optical action potential traces acquired from two locations on the epicardial surface (blue and red traces correspond to circled regions of interest on the heart surface shown in panel C). **C)** Time snapshots of epicardial fluorescent voltage images, which show the progression of electrical activation. **D)** Scatter plot comparing the duration time of the uncorrected QT, corrected QT using various formulae, and optically measured QT (QTc_op) values in the same, excised heart preparations (n=44 total). **E)** Comparison of the duration difference (ΔQT) between the QTc_op value and QT values, using various formulae. Wherein ΔQT is the duration difference between QTc_op and the uncorrected QT value, ΔQTcAd is the duration difference between QTc_op and the QTcAd value, etc. Values reported as mean ± SD, with individual replicates shown. Statistical significance determined by 1-way ANOVA with Holm-Sidak correction (****p<0.0001). AUF: Arbitrary unit of fluorescent, QTc_op: optically measured QT, QTcAd: Adaptive QTc, QTcB: Bazett QTc, QTcFram: Framingham QTc, QTcF: Fridericia QTc, QTcH: Hodges QTc, QTcW: Van de Water QTc, Uncorrected QT.

### Comparison of the adaptive formula vs Benatar (age-specific) formula

To the best of our knowledge, the Benatar formula (QTcBe) is the only proposed age-specific QT correction equation^24^. The QTcBe formula successfully mitigated the inverse correlation between QT and RR interval, indicated by nearly horizontal regression lines and negligible R^2^ values (**Figure 5A**). However, when compared to the uncorrected QT values, it is evident that the QTcBe formula significantly increases the predicted QT duration across all individual age groups (**Figure 9B,C**). Notably, in the 1-week age group, the uncorrected QT measured 263.8±14.8 ms, which was extended to 422.5±7.3 ms (p<0.0001) using the QTcBe methodology. Such an exaggeration in the predicted QT interval was not observed with the QTcAd formula.

**Figure 5.**
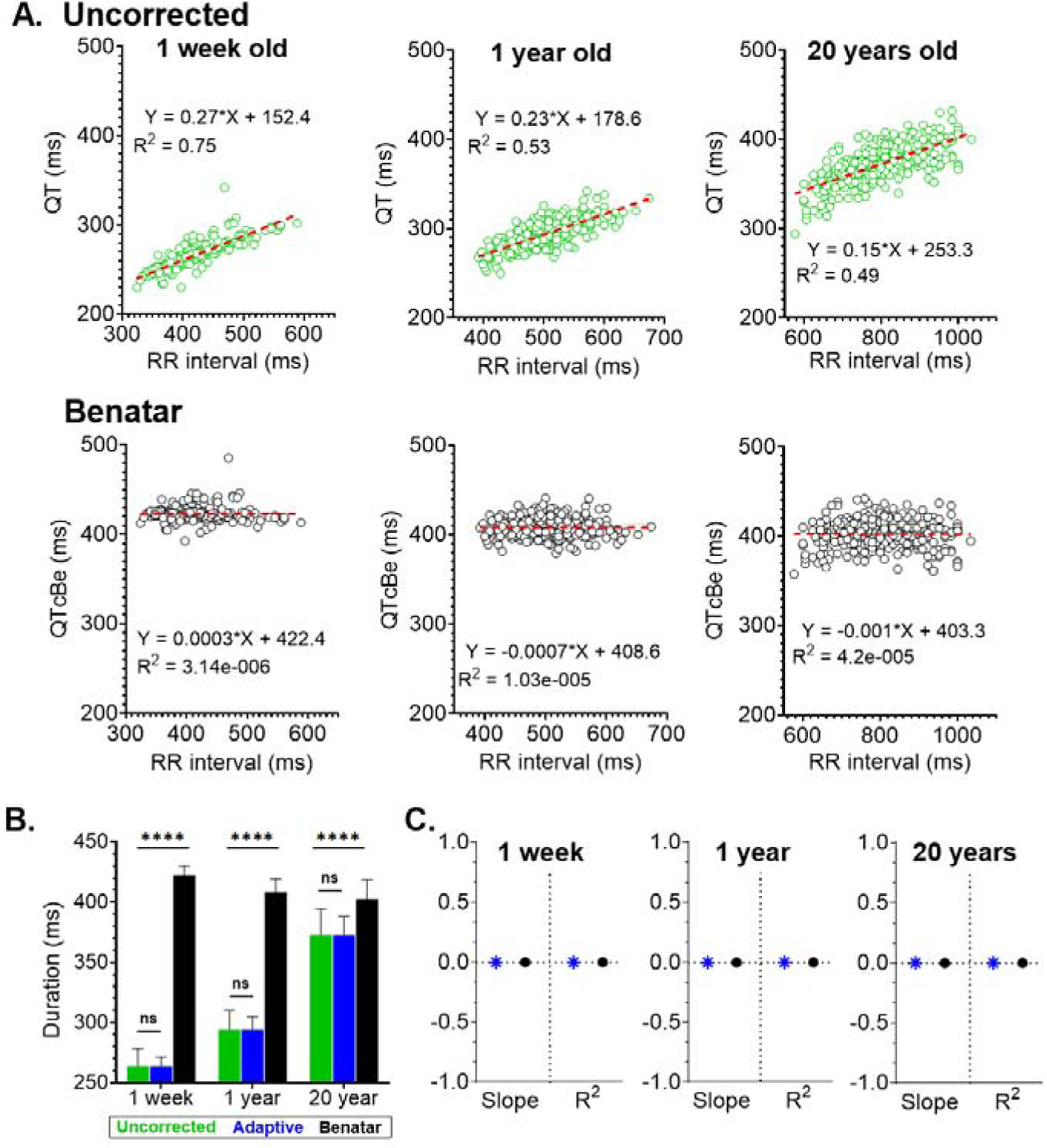
Comparison of adaptive vs Benatar QT (age-specific) correction. Comparison of linear QT-RR interval regression slopes and goodness of fit (R^2^) values derived from human datasets, binned into 1-week (n=300), 1-year (n=300), and 20-years (n=300) age groups. **A)** Top: Uncorrected QT-RR, Bottom: QTcBe-RR. **B)** Comparison of QT duration measurements derived from the uncorrected QT, or using the QTcAd or QTcBe formula. Values reported as mean ± SD, with individual replicates shown. Statistical significance determined by 2-way ANOVA with Holm-Sidak correction (****p<0.0001). Note: exaggerated lengthening of QT value with QTcBe formula, particularly in the younger age groups. **C)** Comparison of QTc-RR linear regression slopes and goodness of fit (R^2^) values from the same cohort. QT: Uncorrected QT, QTcAd: Adaptive QTc, QTcBe: Benatar QTc.

### Influence of QT correction formula on data interpretation

QTc serves as a widely recognized CVD biomarker ^5^ that is crucial for screening the safety and efficacy of anti-arrhythmic medications^27^. To underscore the significance of selecting an appropriate QT correction formula, we investigated whether different formulas impact data interpretation. As shown in **Figure 6** and **Supplemental Figure 4**, the conclusions regarding QTc prolongation or shortening with age varied depending on the correction formula used. For example, in humans, the QTcAd formula shows significant QTc prolongation with advanced age (**Figure 6A**, *1 week*: 263.8±7.4 ms, *1 year*: 294.2±10.7 ms, *20 years*: 372.4±15.5 ms). The QTcB formula showed a modest lengthening of QTc with age (*1 week*: 411.8±11.6 ms, *1 year*: 415.1±15.8 ms, *20 years*: 418±20.1 ms). While both QTcAd and QTcB indicate QTc prolongation with advanced age, data derived using the QTcH formula showed no difference between 1 year and 20 year old individuals (*1 year*: 399.5±14.3 ms, *20 years*: 399.5±19.4 ms). Similar trends of conflicting QTc variation with age were observed in guinea pigs across (**Figure 6B, C**). For instance, *in vivo* ECG recordings in guinea pigs had a longer QTc in older adult animals (203.6±12.2 ms) compared to neonates (182±9.8 ms) when the QTcAd formula was used (**Figure 6B**). While QTcB showed no significant QTc lengthening between neonates and older adults, and QTcH showed significant QTc shortening (*neonate*: 520.7±34.4 ms, *young adult*: 489.3±38 ms, *older adult*: 462.3±43 ms).

**Figure 6.**
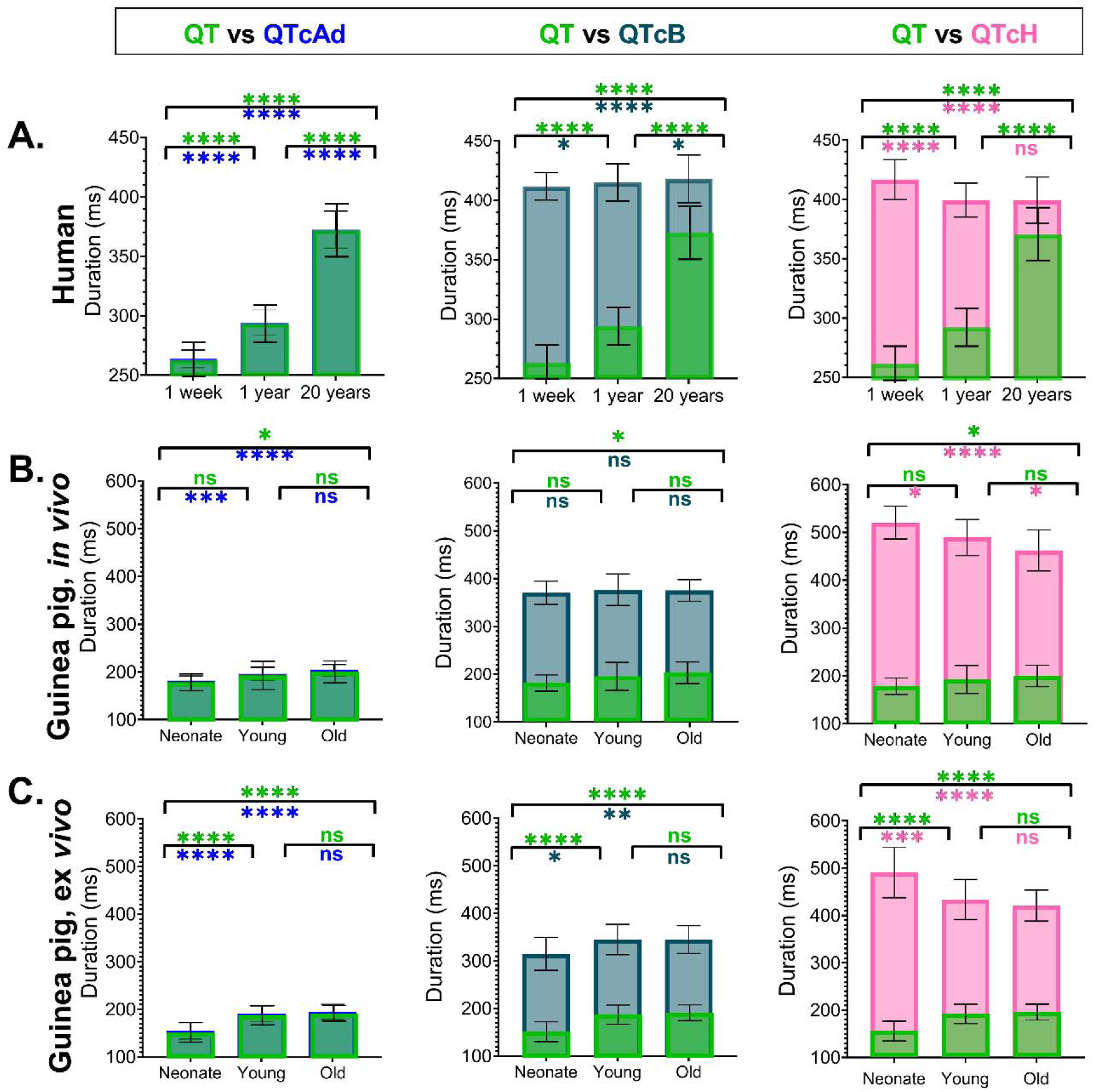
Effect of QTc formula choice on data interpretation. **A)** Comparison of QTcAd, QTcB, and QTcH in three human age groups: 1-week (n=300), 1-year (n=300), and 20-years (n=300). **B)** Comparison of QTcAd, QTcB, and QTcH in guinea pigs (*in vivo*) across three age groups: neonate (n=19), young adult (n=13), and old adult (n=26). **C)** Comparison of QTcAd, QTcB, and QTcH in guinea pigs (*ex vivo*) across three age groups: neonate (n=21), young adult (n=19), and old adult (n=15). Values reported as mean ± SD. The mean uncorrected QT value (green) is superimposed on the mean value derived using the adaptive formula (blue), Bazett formula (teal), or Hodges formula (pink) for each species/age group. Statistical significance determined by 1-way ANOVA with Holm-Sidak correction (*p<0.05, **p<0.01, ***p<0.005, ****p<0.0001).

### Population-based vs individual-specific QT correction

Recent studies have suggested that population-based QT correction methods are inferior due to significant inter-subject variability within a given population, and that individual-specific QT correction methods may be a more accurate alternative^9,20^. Accordingly, we assessed whether the QTcAd formula (population-based) was inferior to an individual-specific correction approach – with the later utilizing a heart rate modification protocol in guinea pigs using a pharmacological agent (ivabradine to slow heart rate) and external atrial pacing (to hasten heart rate). Multiple ECG recordings were collected from each individual heart (n=8) across a range of heart rates, enabling us to model individual ECG datasets (n=149 total ECG datasets). Slopes of the HR-QT linear regression fit varied between animals, ranging from -0.31 to -0.59 (**Figure 7A**). To generate a population-based dataset, the mean resting HR and predicted QT (from linear regression fit) were calculated. From this dataset, the population-based HR-QT regression slope was 0.419 – which closely approximated the mean slope (0.42) of the individual specific HR-QT regression lines. Finally, we generated a testing dataset of QT and HR pairs by randomly selecting 10 data points from each heart preparation, and then we compared values derived from the individual-specific vs a population-based QTcAd formula. As shown in **Figure 7B**, the QTc values were comparable (p=0.397) when derived from either the individual-specific or population-based QTcAd formula.

**Figure 7.**
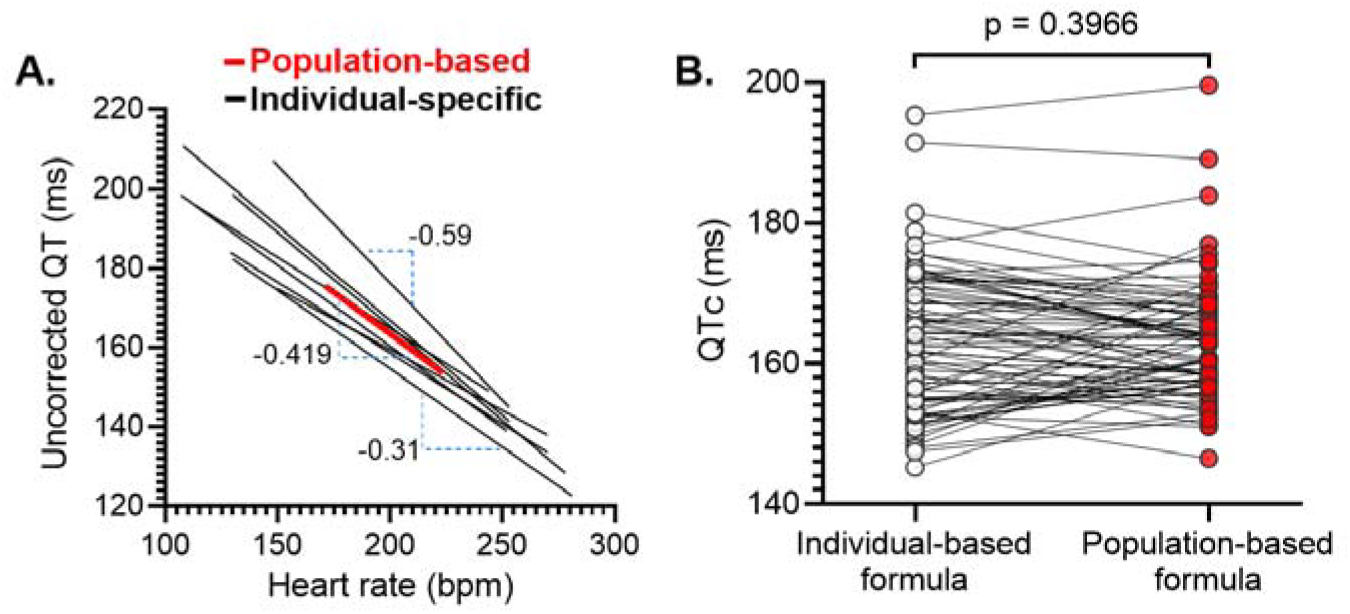
Comparison of population-based vs individual-specific QTc approach. **A)** Linear regression lines depicting individual-specific (black lines) and population-based (red line) uncorrected QT values, using an *ex vivo* ECG dataset obtained from excised, guinea pig hearts subjected to a heart rate adaptation protocol. A total of n=149 total ECG recordings were used. The mean individual-specific regression slope ranged from -0.31 to -0.59, while the mean population-based regression slope was 0.419. **B)** Comparison of QTc values derived from the application of the QTcAd formula using both individual-specific and population-based datasets acquired from the same animals. There was no significant difference between the two approaches (paired, non-parametric Wilcoxon test). Individual replicates are shown.

## Discussion

In this study, we have developed and validated an adaptive QTc formula that incorporates age as a demographic variable. This proposed formula can be applied in two distinct contexts. Equation 1 represents its utilization in research studies, where a dataset containing QT and HR values for the study population is easily accessible. Alternatively, Equation 2 provides a practical application for clinical settings, allowing the prediction of a patient’s QTc by inputting their age, QT, and HR values into the formula. Further discussion below delves into the performance and validation of this adaptive formula.

### Adaptive formula is superior to other commonly used QT correlation formulae

Since the introduction of the Bazett formula, over 20 QT correction formulae have been proposed for study populations with distinct demographics^10^. However, none of these formulae have provided appropriate correction, leaving researchers and clinicians searching for an ideal QT correction formula ^6,8–10,18^. Multiple studies have shown that correction formulae, including Bazett, may demonstrate robustness in one study population but fail in another ^6,14,18,19,28^. This observation led us to hypothesize that animal subjects or patient populations with unique demographics likely require a specific correction formula, which can be tailored according to the demographic characteristics of the cohort.

Among the widely used formulae, Bazett and Fredericia corrections are non-linear and rely on the instantaneous RR interval, making it challenging to adapt these formulae to varying QT-HR relationships in diverse populations. Conversely, formulae such as Hodges and Framingham offer linear correction and are based on the instantaneous HR or RR interval (respectively), along with corresponding linear regression coefficients. However, as suggested in the current study, the accuracy of these formulae is limited without proper adjustment. Accordingly, we proposed a modified linear-correlation formula that takes into account and utilizes the linear regression slopes and mean resting HR of a specific population with unique demographics. In this study, as proof of concept, we used animal or patient age as the main demographic variable. Our results demonstrate that this adaptive approach overcomes the traditional limitations associated with linear correction, and the proposed formula is superior to other widely used formulae (**Figure 3**).

### Adaptive formula offers appropriate age-specific QT correction

Age is a crucial confounding demographic variable that significantly affects the electrical properties of the heart ^3,4,24,25,29–31^. Previous studies have shown a linear and inverse correlation between age-specific HR and the QT interval^18^. However, both HR and the QT interval are influenced by age in both human ^1,2^ and animal models^3,4^. Our findings align with these previous studies, demonstrating significant age-dependent changes in HR and QT (see **Supplemental Table 1 and 2**). These findings led us to hypothesize that QT correction formulae require “adaptation” to take into account variations in the linear regression slope and mean resting HR within a specific age group (see **Table 1**). To the best of our knowledge, the QTcBe formula is the only age-specific QT correction formula available. As such, we compared the effectiveness of our adaptive QTcAd formula to QTcBe. Despite slight differences in the human age groups used in Benatar, et al.^24^ and our current study, the uncorrected QT-RR regression slopes were comparable (Benatar, et al: *0-1 year m:* 0.24, *1-5 years m:* 0.24, *>10 years m:* 0.18; Current study: *1 week m:* 0.27, *1 year m:* 0.23, *20 years m:* 0.15). QTcBe successfully eliminated the linear correlation between QT and the RR interval (**Figure 5A**), but, it also led to a significant QTc prolongation – particularly at faster heart rates – which did not occur with the QTcAd formula. A thorough investigation of the observed underperformance of QTcBe in younger age groups is beyond the scope of this study. However, one plausible explanation could be the use of an instantaneous RR interval instead of a mean or median value in the formula. Further, the linearity assumption between QT-RR correlation used to develop the QTcBe formula has been debated^18^.

### The predicted QT interval is more accurately approximated by the QTcAd formula compared to other formulae

In this study, we experimentally validated the QTcAd formula-derived QT approximation of ventricular activation-repolarization time during sinus rhythm. Additionally, we demonstrated that compared to other corrected QT formulae, QTcAd provides a more accurate approximation of activation-repolarization time (**Figure 4D,E**). Importantly, our results demonstrate that QT correction is frequently overestimated by commonly employed formulae tested in this study. To the best of our knowledge, this is the first study to validate QT correction using an experimental optical mapping approach. Notably, the QT time approximated from optical voltage signals is shorter than the QT interval measured from *ex vivo* pseudo-ECG recordings. This difference is explained by the fact that QTc_op is measured by assuming that activation time corresponds to breakthrough time – but, it does not account for the conduction delay from the AV node to the earliest epicardial breakthrough site.

### Choice of QT correction formula influences data interpretation

Our results suggest that the choice of QT correction formula significantly influences data interpretation and conclusions – which can significantly impact studies with demographic heterogeneity and/or drug safety assessments. As shown in **Figure 6** and **Supplemental Figure 4**, statistically significant differences in QTc values across age groups varied depending on the correction formula applied. For example, the QTcAd formula suggested QTc prolongation occurs with advanced age (in both humans and guinea pigs) – while QTcH suggested QTc shortening. We also observed that many of the widely used correction formulae exaggerated the predicted QTc interval time – as compared to the raw uncorrected values (**Figure 5**, **7**). This observation that QT correction formula choice significantly influences data interpretation is in agreement with other reports – include those that tested different QT correction methods for long QT syndrome screening in both human subjects and animal models ^17,28,32,33^. For example, one study reported that QTcB was associated with more false positives in pediatric patients compared to either QTcF or QTcFram^32^. While another study demonstrated that QTcB was a superior diagnostic tool, as compared to either QTcF or QTcFram^17^. With respect to age-dependent adaptations in the QT interval, Benatar et al.^24^ demonstrated that use of QTcFram suggested a significant age-dependent QTc prolongation in male children (+36 ms), QTcB suggested a non-significant prolongation (+8ms), and QTcBe suggested no prolongation. However, Rijnbeek et al. ^34^ used a similar age-group of male children and reported a non-significant shortening of QTc with age (0-1 month vs 8-12 years, -2ms). Such inconsistencies are also widely observed in animal models. For example, use of the QTcFram formula showed a significant QTc prolongation with advanced age – yet, in our prior study, we did not detect such a robust age-dependent difference when QTcB was applied^35^.

### Population-based QTc formula is not inferior to an individual-specific formula

QTc correction formulae are widely used in clinics and experimental research studies, however, the accuracy of using a universal formula has been disputed in both human subjects and animal models^9,20^. Indeed, it has been suggested that an individual-specific QT correction formula may be a superior approach, as it mitigates inter-subject variability that is observed in ECG parameters. Contrary to these findings^9,20^, our experimental study compared QTc values in guinea pigs using both individual-specific and population-based datasets, which resulted in comparable values when the QTcAd formula was applied (**Figure 7**). Nevertheless, the aforementioned studies^9,20^ correctly identified a major weakness in using a universal one-size-fits all QT correction formula - as demographic variables significantly modify ECG characteristics. This limitation has also been highlighted by other studies^25,36,37^. Importantly, widely used population-based QT correction formulae were all developed using ECG datasets from individuals with variable age ranges (e.g., Bazett: 14-53 years; Framingham: 28-62 years; Hodges: 20-89 years; Fridericia: 30-81 years) – which could impact the formula parameters and may explain conflicting results on QT correction performance^6,10^. Indeed, a study by Vandenberk et al.^6^ demonstrated that in a population with an average age of 59.8±16.2 years - QTcF and QTcFram formulae (developed based on datasets from an older population) outperformed QTcB (developed based on a younger population). Accordingly, the QTcAd formula can serve as a surrogate, as the specified parameters can be adjusted according to the demographic characteristics of a specific population.

## Limitations

This study proposed and validated an adaptive QT correction formula, focused solely on age as a demographic variable. Although age is a significant factor, other variables such as sex and race, can also influence demographic characteristics^38^. The current QTcAd formula does not account for these variables, which may require additional modifications to create a comprehensive "demography"-based formula. The effectiveness of the QTcAd formula was tested using both human and guinea pig ECG datasets and validated through experimental methods. However, the efficacy of using the QTcAd formula for diagnostic capabilities or therapeutic effects was not assessed and will require additional future investigations. In this study, the dataset used to develop the QTcAd formula was restricted to a young human population (1360 individuals), which neglects additional changes to QT-HR dynamics that can occur in later adulthood. Expanding the human dataset to include older individuals would likely enhance the overall utility of the QTcAd formula across a broader age range. We also provided an estimation of the regression slope and resting basal heart rate, using eight age groups. However, considering the rapid dynamic changes in cardiac electrophysiology early in life, age-specific correction may benefit from a larger dataset that spans more age groups to improve its predictability. Finally, we evaluated both individual-specific and population-based forms of the QTcAd formula in guinea pigs – a similar evaluation in humans would require a clinical trial to adjust each individual patient’s heart rate, which was beyond the scope of this study.

## Conclusions

Our study highlighted the superiority of the proposed QTcAd formula, which incorporated age as a demographic variable, over several commonly used universal QTc formulae. Experimental validation indicated that the predicted QTcAd value closely approximated the activation-repolarization time in an isolated, intact heart model. Furthermore, our findings demonstrated that the values derived from the QTcAd formula is more accurate than the age-specific QTcBe formula. Additionally, the application of the QTcAd formula suggested that a population-based formula is not inferior to individual-specific QTc formulae.

## Abbreviations

BPM: Beats per minute
ECG: Electrocardiogram
HR: Heart Rate
QTc: Corrected QT
QTcAd: Adaptive QTc
QTcB: Bazett QTc
QTcBe: Benatar QTc
QTcFram: Framingham QTc
QTcF: Fridericia QTc
QTcH: Hodges QTc
QTc_op: Optically approximated QT
QTcW: Van de Water QTc

## Sources of Funding

This work was supported by the National Institutes of Health grants R01HD108839 (NGP), Children’s National Research Institute, Children’s National Heart Institute, and Sheikh Zayed Institute for Pediatric and Surgical Innovation.

## Acknowledgements

We gratefully acknowledge Shatha Salameh, Luther Swift, Anika Haski, Anysja Roberts, and Blake Cooper for technical assistance related to ECG recordings.

## Conflict of Interest

None declared.

## Data Availability

Derived data supporting the findings of this study are available from the corresponding author (NGP) upon request.

## Notes

### Competing Interest Statement

The authors have declared no competing interest.

