## Supplemental File for "Validation of a Demography-Based Adaptive QTc Formula using Pediatric and Adult Datasets Acquired from Humans and Guinea Pigs"

**Supplemental Methods and Figures**

Corrected QT formula derivation

For a dataset Ω consisting of ordered pairs {(*x_1_*, *y_1_*), (*x_2_*, *y_2_*), …..., (*x_n_*,*y_n_*)} the linear regression fit is given by,

$\hat{y}=mx+c$ ………………………………… eq.1,

where the y-intercept is given by, $c=\bar{y}-m\bar{x}$, where $\bar{y}=\frac{\sum y}{n}$, $\bar{x}=\frac{\sum x}{n}$, and *m=* slope of the line. From eq.1,

$\hat{y}=\bar{y}+m(x-\bar{x})$

The residual between the observed values (*y*) and predicated $(\hat{y})$ values of the linear fit can be given by,

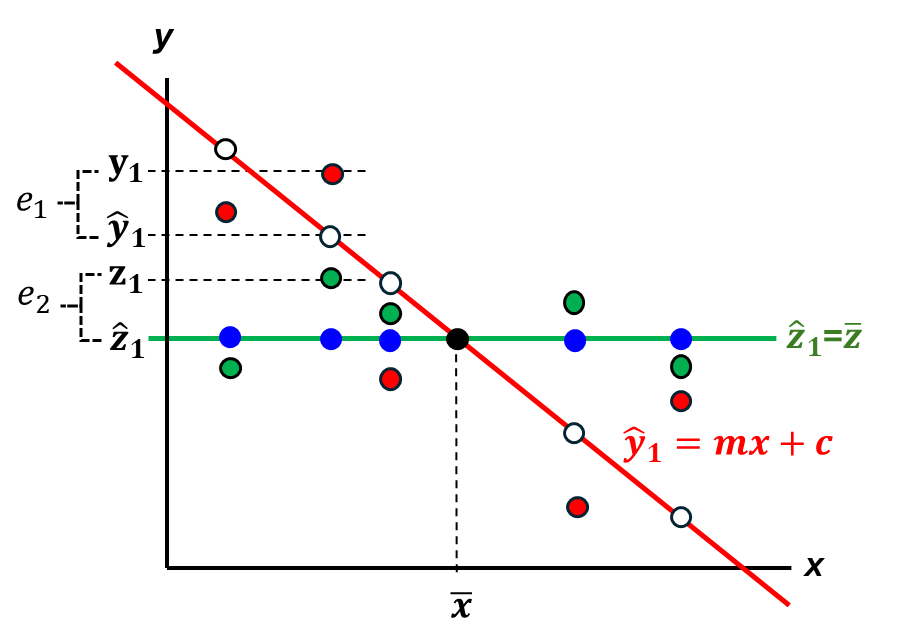
 $e_{1}=y-\hat{y}=y-\bar{y}-m(x-\bar{x})$…………… eq.2

**Supplemental Figure 1: Derivation of QTc (projected) from QT (observed).** Red line is the regression fit of the observed dataset (Ω, red circles). Green line is the regression fit of the projected dataset (ℬ, green circles) of the observed dataset Ω. *e_1_* is the residual between the observed values (*y*) and predicated $(\hat{y})$ values of the linear fit of the dataset Ω. *e_2_* is the residual between the projected values (z) and predicted values ($\hat{z}$) of the linear fit of the projected dataset ℬ.

Let's consider another dataset, ℬ, represented by ordered pairs {(*x_1_*, *z_1_*), (*x_2_*, *z_2_*), …..., (*x_n_*,*z_n_*)} , which is a projection of dataset Ω. The linear fit for dataset ℬ is given by:

$\hat{z}=m_{1}z+c_{1}$…………………………………eq.3

where the y-intercept is given by, $c_{1}=\bar{z}-m_{1}\bar{x}$, where $\bar{z}=\frac{\sum z}{n}$, $\bar{x}=\frac{\sum x}{n}$, and $m_{1}$is the slope of the fitted line. To achieve a null correlation ($R^{2}\cong0)$ between the projected values (z) and the predicted values ($\hat{z}$) eq.3 must satisfy the condition $m_{1}$= 0. Therefore, y- intercept $c_{1}=\bar{z}$ , and from eq.3,

$\hat{z}=\bar{z}$ ………………………………………. eq.4

Since, fitted line of dataset Ω intersects fitted line of dataset ℬ at $\bar{z}$,

$\hat{z}=\bar{z}=\bar{y}$

The residual between the projected values (z) and predicted values ($\hat{z}$) of the linear fit can be given by,

$e_{2}=z-\overline{z}= z-\overline{y}$ …………………… eq.5

Since the regression line of dataset ℬ represents the mean of the observed values (the mean line), and the regression line of dataset Ω intercepts the mean line, it follows that the two regression models will have equal residual values for each point. Therefore,

$e_{1}=e_{2}$

Thus, from eq.2 and eq.5,

$z=y-m(x-\overline{x})$ ………………………. eq.6

In cases where the independent variable (x) is inversely correlated with the dependent variables (*y, z*) the slope of the regression line is always negative, and eq.6 becomes,

$z=y+m(x-\overline{x})$ ……………………… eq.7

Substituting the variables in equation 7, where *z* is replaced by corrected QT (QT_c_), *y* is replaced by QT, x is replaced by heart rate (HR) and $\overline{x}$ is replaced by the mean of the heart rate ($\bar{\mathrm{HR}}$), yields:

QT_c_ = QT+ |*m|*(HR-$\bar{\mathrm{HR}}$) ……………………. eq.8

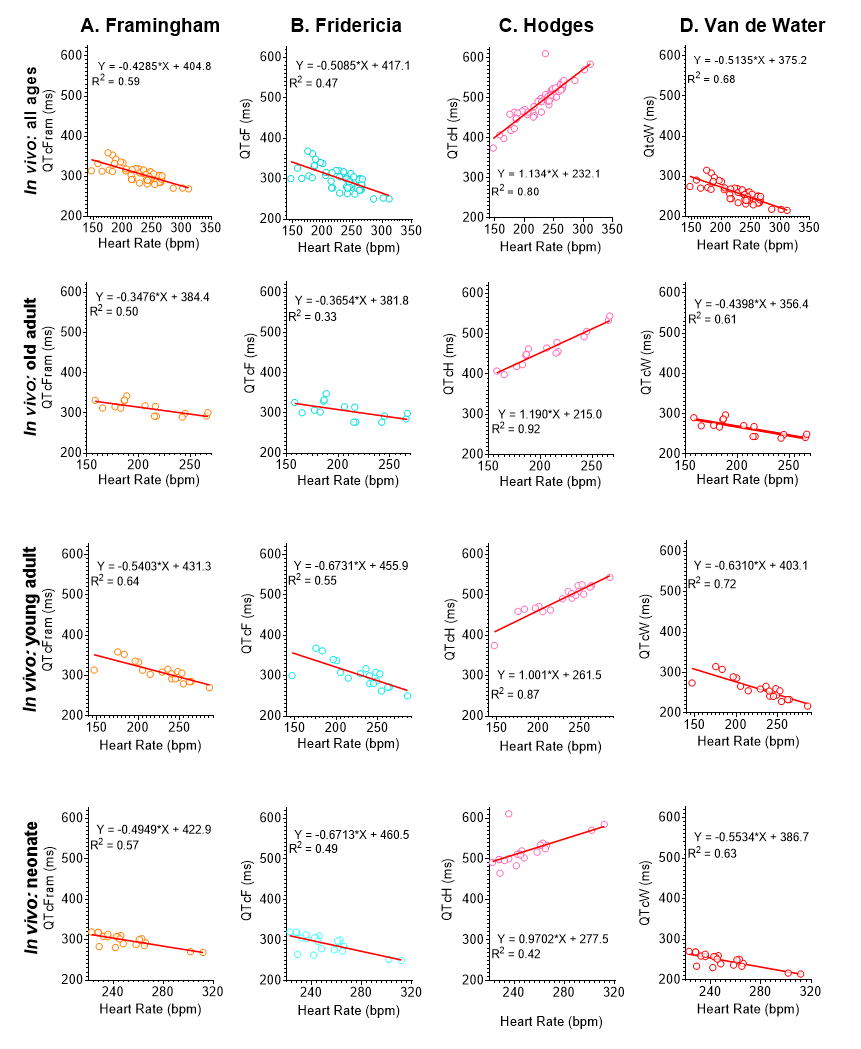

**Supplemental Figure 2. Comparison of QTc formulae across specific guinea pig age groups.** *Ex vivo* linear regression analysis of the QT-HR relationship in guinea pigs across 4 different age groups, including all ages, neonates (n=21), young adults (n=19), and old adults (n=15). **A)** QTcFram-HR, **B)** QTcF-HR, **C)** QTcH-HR, **D)** QTcW-HR derived from the same animals and age groups. Corresponding equations and goodness of fit (R2) are provided. QTcFram: Framingham QTc, QTcF: Fridericia QTc, QTcH: Hodges QTc, QTcW: Van de Water QTc, bpm = beats per minute.

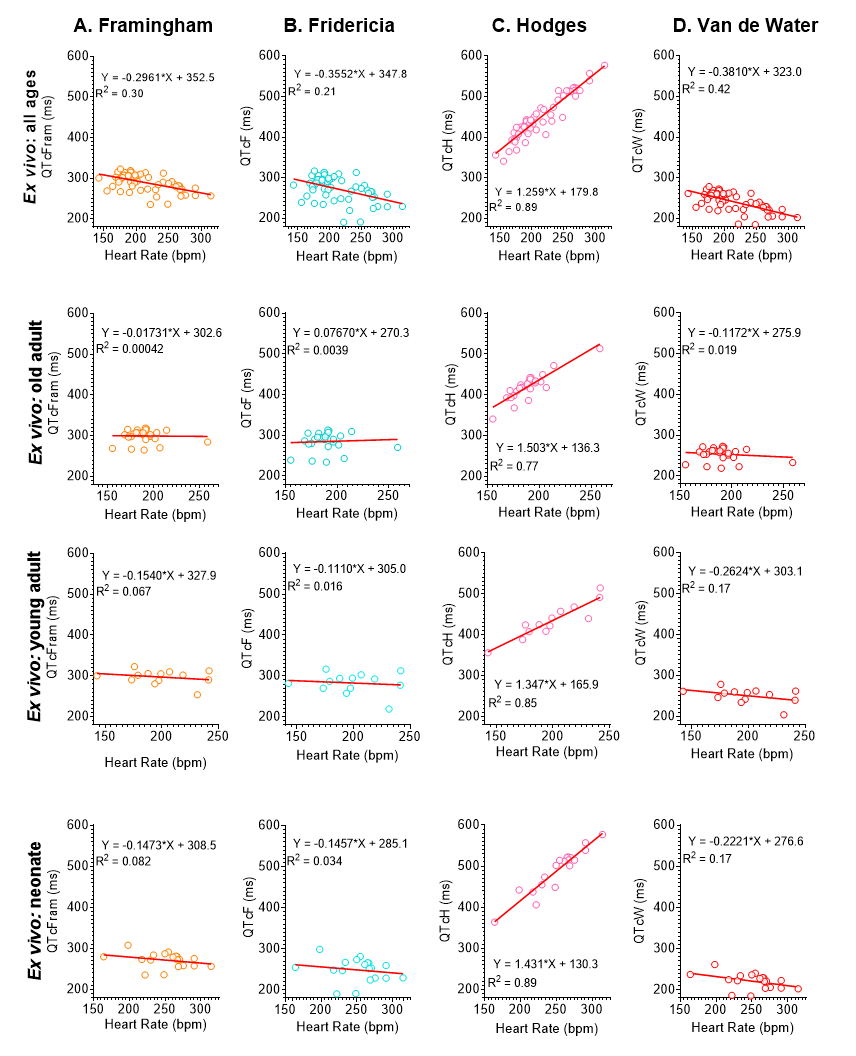

**Supplemental Figure 3. Comparison of QTc formulae across specific guinea pig age groups.** *Ex vivo* linear regression analysis of the QT-HR relationship in guinea pigs across 4 different age groups, including all ages, neonates (n=19), young adults (n=13), and old adults (n=26). **A)** QTcFram-HR, **B)** QTcF-HR, **C)** QTcH-HR, **D)** QTcW-HR derived from the same animals and age groups. Corresponding equations and goodness of fit (R2) are provided. QTcFram: Framingham QTc, QTcF: Fridericia QTc, QTcH: Hodges QTc, QTcW: Van de Water QTc, bpm = beats per minute.

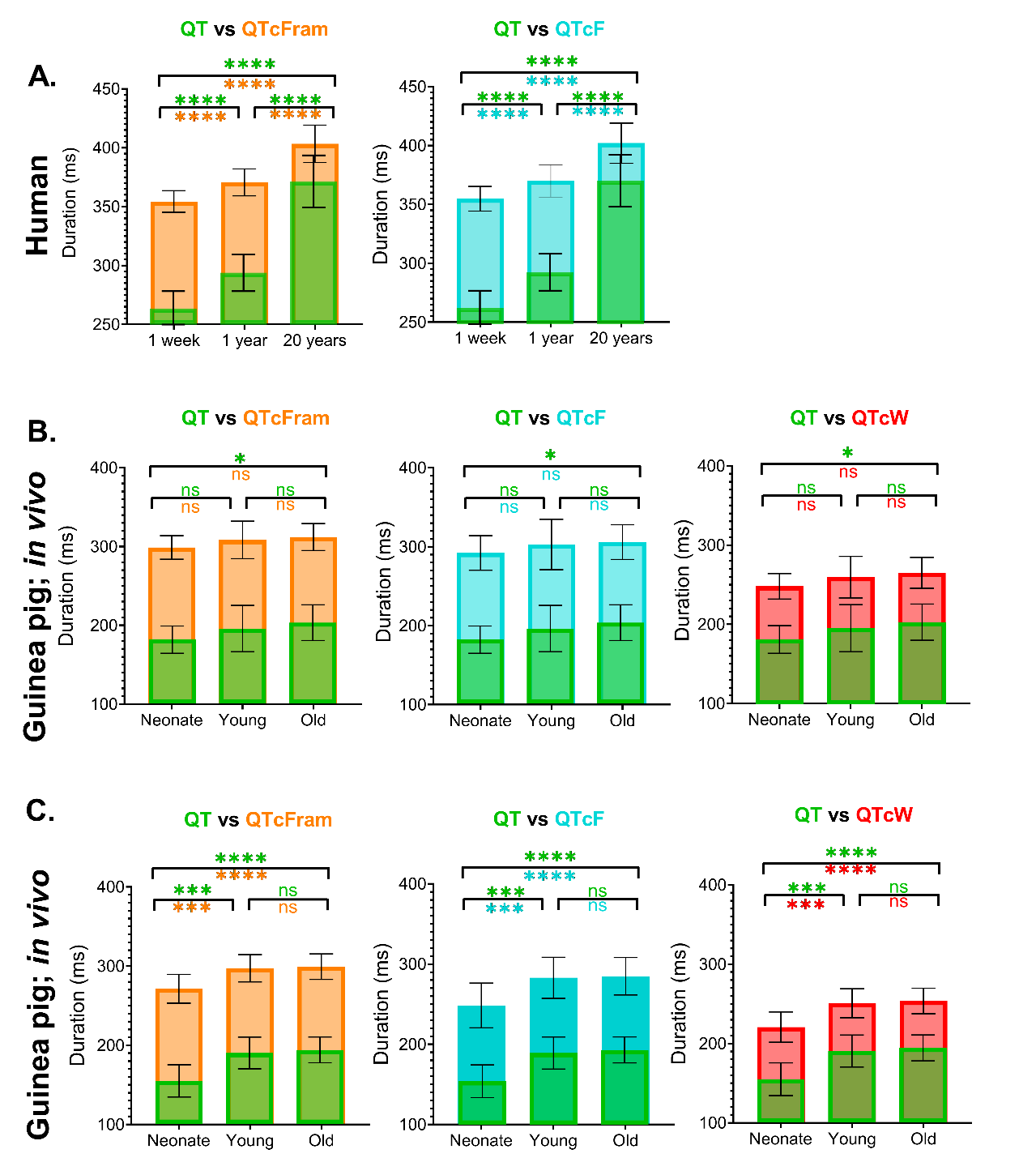

**Supplemental Figure 4. Effect of QTc formula on data interpretation**. **A)** Comparison of uncorrected QT, QTcFram, and QTcF in three human age groups: 1-week (n=300), 1-year (n=300), and 20-years (n=300). **B)** Comparison of uncorrected QT, QTcFram, QTcF, and QTcW in guinea pigs (*in vivo*) across three age groups: neonate (n=19), young adult (n=13), and old adult (n=26). **C)** Comparison of uncorrected QT, QTcFram, QTcF, and QTcW in guinea pigs (*ex vivo*) across three age groups: neonate (n=21), young adult (n=19), and old adult (n=15). Values reported as mean ± SD. The mean uncorrected QT value (green) is superimposed on the mean value derived using the Framingham formula (orange), Fridericia formula (cyan), or Van de Water formula (red) for each species/age group. Statistical significance determined by 1-way ANOVA with Holm-Sidak correction (*p<0.05, **p<0.01, ***p<0.005, ****p<0.0001).

**Supplemental Tables**

**Supplemental Table 1. Guinea pig demographic and ECG characteristics.** Comparison between two groups were performed using a Student’s t-test. Statistical significance denoted *p<0.05, **p<0.005, ****p<0.0001

| **Experiment** | **Age group** | **n** | **HR (bpm)**  **(mean ± SD)** | **QT (ms)**  **(mean ± SD)** | **Comparison** | **HR**  ***p*** | **QT**  ***p*** |
| --- | --- | --- | --- | --- | --- | --- | --- |
| *In-vivo*  (CNRI) | Neonate (1-2 days) | 19 | 251.8 ± 35.2 | 154.8 ± 35.2 | Neonate vs. Young | * | ns |
|  | Younger adult (1-3 months) | 13 | 199.1 ± 28.8 | 190.6 ± 20.2 | Neonate vs. Old | *** | * |
|  | Older adult (18-22 months) | 26 | 189.4 ± 19.0 | 194.4 ± 16.4 | Young vs. Old | ns | ns |
| *Ex-vivo*  Langendorff-heart preparation (CNRI) | Neonate (1-2 days) | 21 | 250.7 ± 22.9 | 181.9 ± 17.4 | Neonate vs. Young | **** | **** |
|  | Younger adult (1-3 months) | 19 | 227.5 ± 35.2 | 196.1 ± 29.6 | Neonate vs. Old | **** | **** |
|  | Older adult (18-22 months) | 15 | 207.7 ± 34.6 | 203.7 ± 22.8 | Young vs. Old | ns | ns |
| *Ex-vivo*  Working heart preparation (KCL) | Younger adult (300-350 g) | 8 | 258 ± 14.1 | 139.5 ± 5.9 | - | - | - |

**Supplemental Table 2. Human subject demographic and ECG characteristics.** Comparison between two groups were performed using a Student’s t-test. Statistical significance denoted *p<0.05

| **Age** | **Sex** | **n** | **HR (bpm)**  **(mean ± SD)** | **QT (ms)**  **(mean ± SD)** | **Heart Rate**  ***p* value**  **(sex)** | **QT**  ***p* value**  **(sex)** |
| --- | --- | --- | --- | --- | --- | --- |
| 1 week | Male | 149 | 147 ± 16.1 | 263 ± 14.2 | ns | ns |
|  | Female | 151 | 148 ± 15.9 | 264 ± 15.4 |  |  |
| 1 year | Male | 160 | 120 ± 11.8 | 294 ± 16 | ns | ns |
|  | Female | 140 | 120 ± 12.2 | 295 ± 15.4 |  |  |
| 2 years | Male | 50 | 112 ± 15 | 302 ± 21.7 | ns | ns |
|  | Female | 50 | 114 ± 12.1 | 300 ± 19.6 |  |  |
| 5 years | Male | 59 | 96 ± 12.2 | 328 ± 19.9 | ns | ns |
|  | Female | 41 | 95.2 ± 14.7 | 330 ± 25.5 |  |  |
| 10 years | Male | 57 | 81 ± 13.9 | 360 ± 26.7 | ns | * |
|  | Female | 43 | 86 ± 13.1 | 350 ± 25.1 |  |  |
| 15 years | Male | 28 | 74 ± 10.5 | 370 ± 24.2 | ns | ns |
|  | Female | 72 | 75.8 ± 10.6 | 369 ± 22 |  |  |
| 20 years | Male | 89 | 75 ± 10.5 | 368 ± 20 | ns | * |
|  | Female | 211 | 76.8 ± 9.81 | 374 ± 22.8 |  |  |
| 25 years | Male | 10 | 70.8 ± 10.7 | 385 ± 22.1 | ns | ns |
|  | Female | 50 | 76.7 ± 11.4 | 380 ± 25.2 |  |  |

**Supplemental Table 3. QTc-HR regression equation and R^2^ values in Humans**

| **Formula** | **QT** | **Bazett** | **Fridericia** | **Framingham** | **Hodges** | **Adaptive** |
| --- | --- | --- | --- | --- | --- | --- |
| 1 week | y = -0.80x + 382.17  R² = 0.75 | y = 0.19x + 383.43 R² = 0.071 | y = -0.25x + 391.13 R² = 0.14 | y = -0.35x + 405.9 R² = 0.37 | y = 0.95x + 277.17 R² = 0.81 | y = 2E-05x + 263.81 R² = 2E-09 |
| 1 year | y = -0.96x + 409.72  R² = 0.54 | y = 0.35x + 373.44 R² = 0.070 | y = -0.20x + 393.83 R² = 0.030 | y = -0.33x + 409.7 R² = 0.12 | y = 0.79x + 304.72 R² = 0.44 | y = -0.0009x + 294.32 R² = 9E-07 |
| 20 years | y = -1.58x + 492.75  R² = 0.51 | y = 0.91x + 348.46 R² = 0.21 | y = 0.013x + 400.99 R² = 6E-05 | y = -0.0013x + 403.37 R² = 7E-07 | y = 0.17x + 387.75 R² = 0.012 | y = -3E-05x + 372.44 R² = 5E-10 |
| All Ages | y = -1.46x + 477.81  R² = 0.91 | y = -0.0094x + 416.01 R² = 0.00030 | y = -0.59x + 443.25 R² = 0.61 | y = -0.63x + 447.74 R² = 0.70 | y = 0.29x + 372.81 R² = 0.28 | y = -3E-05x + 310.2 R² = 5E-09 |

**Supplemental Table 4. QTc-HR regression equation and R^2^ values in guinea pigs (in-vivo)**

| **Formula** | **QT** | **Bazett** | **Fridericia** | **Framingham** | **Hodges** | **Van De Water** | **Adaptive** |
| --- | --- | --- | --- | --- | --- | --- | --- |
| Neonate | y = -0.63x + 339.77  R² = 0.68 | y = -0.63x + 528.46 R² = 0.35 | y = -0.67x + 460.5 R² = 0.49 | y = -0.49x + 422.92 R² = 0.57 | y = 0.97x + 277.48 R² = 0.42 | y = -0.55x + 386.75 R² = 0.63 | y = -2E-05x + 181.98 R² = 2E-09 |
| Young | y = -0.75x + 366.52  R² = 79 | y = -0.55x + 500.62 R² = 0.34 | y = -0.67x + 455.86 R² = 0.55 | y = -0.54x + 431.26 R² = 0.64 | y = 1.00x + 261.52 R² = 0.87 | y = -0.63x + 403.1 R² = 0.72 | y = -3E-05x + 196.11 R² = 5E-09 |
| Old | y = -0.56x + 319.99  R² = 0.72 | y = -0.15x + 407.07 R² = 0.055 | y = -0.37x + 381.8 R² = 0.33 | y = -0.35x + 384.36 R² = 0.50 | y = 1.19x + 214.99 R² = 0.92 | y = -0.44x + 356.35 R² = 0.61 | y = 2E-05x + 203.59 R² = 2E-09 |
| All Ages | y = -0.62x + 336.88  R² = 0.76 | y = -0.35x + 455.51 R² = 0.21 | y = -0.51x + 417.08 R² = 0.47 | y = -0.43x + 404.76 R² = 0.59 | y = 1.13x + 232.09 R² = 0.80 | y = -0.51x + 375.23 R² = 0.68 | y = 5E-06x + 192.78 R² = 2E-10 |

**Supplemental Table 5. QTc-HR regression equation and R^2^ values in guinea pigs (ex-vivo)**

| **Formula** | **QT** | **Bazett** | **Fridericia** | **Framingham** | **Hodges** | **Van De Water** | **Adaptive** |
| --- | --- | --- | --- | --- | --- | --- | --- |
| Neonate | y = -0.32x +235.27  R² = 0.30 | y = 0.039x + 305.08 R² = 0.0016 | y = -0.15x + 285.13 R² = 0.034 | y = -0.15x + 308.47 R² = 0.082 | y = 1.43x + 130.27 R² = 0.89 | y = -0.22x + 276.63 R² = 0.17 | y = -2E-05x + 154.87 R² = 2E-09 |
| Young | y = -0.40x + 270.87  R² = 0.33 | y = 0.16x + 313.26 R² = 0.021 | y = -0.11x + 304.96 R² = 0.016 | y = -0.15x + 327.87 R² = 0.067 | y = 1.35x + 165.87 R² = 0.85 | y = -0.26x + 303.07 R² = 0.17 | y = -5E-05x + 190.6 R² = 7E-09 |
| Old | y = -0.25x + 241.28  R² = 0.081 | y = 0.37x + 275.38 R² = 0.057 | y = 0.077x + 270.3 R² = 0.0039 | y = -0.017x + 302.57 R² = 0.00040 | y = 1.50x + 136.28 R² = 0.77 | y = -0.12x + 275.9 R² = 0.019 | y = -9E-06x + 194.59 R² = 1E-10 |
| All Ages | y = -0.49x + 284.78  R² = 0.55 | y = -0.19x + 376.18 R² = 0.049 | y = -0.36x + 347.8 R² = 0.21 | y = -0.30x + 352.49 R² = 0.30 | y = 1.26x + 179.78 R² = 0.89 | y = -0.38x + 323.03 R² = 0.42 | y = 5E-08x + 180.63 R² = 3E-15 |
